# The C-terminal Helix 9 motif regulates cannabinoid receptor type 1 trafficking and surface expression

**DOI:** 10.1101/501528

**Authors:** Alexandra Fletcher-Jones, Keri L. Hildick, Ashley J. Evans, Yasuko Nakamura, Kevin A. Wilkinson, Jeremy M. Henley

## Abstract

Cannabinoid type 1 receptor (CB1R) is only stably surface expressed in axons, where it downregulates neurotransmitter release. How this tightly regulated axonal surface polarity is established and maintained is unclear. To address this question, we used time-resolved imaging to determine the trafficking of CB1R from biosynthesis to mature polarised localisation. We show that the secretory pathway delivery of CB1R is axonally biased and that surface expressed CB1R is more stable in axons than in dendrites. This dual mechanism is mediated by the CB1R C-terminal and involves the Helix 9 (*H9*) domain. Removal of the *H9* domain increases dendrite secretory pathway delivery and decreases in surface stability. Furthermore, CB1R^ΔH9^ is more sensitive to agonist-induced internalisation and less efficient at downstream signalling than CB1R^WT^. Together, these results shed new light on how polarity of CB1R is mediated and indicate that the C-terminal *H9* domain plays key roles in this process.

## Introduction

CB1R is one of the most abundant G-protein-coupled receptors (GPCRs) in the CNS and endocannabinoid signalling through CB1R is a neuromodulatory system that influences a wide range of brain functions including pain, appetite, mood, and memory (Lu and Mackie, 2016; Soltesz et al., 2015). Furthermore, CB1R function and dysfunction are implicated in multiple neurodegenerative disorders (Basavarajappa et al., 2017). Thus, modulation of endocannabinoid pathways is of intense interest as a potential therapeutic target (Reddy, 2017).

CB1R is present in both excitatory and inhibitory neurons, and also in astroglia, where it plays important roles in synaptic plasticity and memory (Busquets-Garcia et al., 2018; Han et al., 2012; Robin et al., 2018). In hippocampal neurons, ~80% of CB1R is present in intracellular vesicular clusters in the soma and dendrites (Leterrier et al., 2006). Strikingly, however, CB1R is not stably surface expressed on somatodendritic plasma membrane. Rather, it has a highly polarised axonal surface expression (Coutts et al., 2001; Irving et al., 2000) where it acts to attenuate neurotransmitter release (Katona, 2009) and modulate synaptic plasticity (Lu and Mackie, 2016).

How this near exclusive axonal surface expression of CB1R is established remains the subject of debate. One suggestion is that high rates of endocytosis due to constitutive activity selectively remove CB1Rs from the somatodendritic compartment, resulting in an accumulation at the axonal surface (Leterrier et al., 2006). These internalised somatodendritic CB1Rs may then be either sorted for degradation or recycled to axons via a transcytotic sorting pathway (Simon et al., 2013). Alternatively, newly synthesized CB1R may be constitutively targeted to lysosomes, but under appropriate circumstances the CB1Rs destined for degradation are retrieved and rerouted to axons (Rozenfeld, 2011; Rozenfeld and Devi, 2008).

Surprisingly, a direct role for the 73-residue intracellular C-terminal domain of CB1R (ctCB1R) in axonal/somatodendritic trafficking or polarised surface expression has not been identified. It has, however, been reported that motifs within ctCB1R are required for receptor desensitization and internalization (Hsieh et al., 1999; Jin et al., 1999) (reviewed by (Mackie, 2008)). Interestingly, there are two putative helical domains in ctCB1R (*H8* and *H9*). *H8* has been proposed to play a role in ER assembly and/or exit during biosynthesis (Ahn et al., 2010; Stadel et al., 2011). The role of the 21-residue *H9* motif is unknown, although analogous regions have been reported to act as a Gαq-binding site in both squid rhodopsin (Murakami and Kouyama, 2008) and bradykinin receptors (Piserchio et al., 2005).

Here we systematically investigated how axonal surface polarity of CB1R arises by tracking newly-synthesised CB1Rs through the secretory pathway to their surface destination. We demonstrate that a population of CB1R is preferentially targeted to the axon through the biosynthetic pathway. CB1Rs that reach the dendritic membrane are rapidly removed by endocytosis whereas CB1Rs surface expressed on the axonal membrane have a longer residence time. We further show that the putative helical domain *H9* in ctCB1R plays a key role in CB1R surface expression and endocytosis in hippocampal neurons. Taken together our data suggest that CB1R polarity is determined, at least in part, by a novel determinant in the C-terminus of CB1R that contributes to targeted delivery to the axonal compartment and the rapid removal of CB1Rs that reach the somatodendritic membrane.

## Results

### Preferential delivery of newly synthesized CB1Rs to, and retention at, the axonal membrane establishes surface polarisation

To investigate how CB1R surface polarity is established we used the retention using selective hooks (RUSH) system (Boncompain et al., 2012) to examine its secretory pathway trafficking. CB1R was tagged at the N-terminus with streptavidin binding peptide (SBP) and EGFP (SBP-EGFP-CB1R). When co-expressed with a Streptavidin-KDEL ‘hook’ that localises to the lumen of the Endoplasmic Reticulum (ER), SBP-EGFP-CB1R is anchored at the ER membrane. The retained SBP-EGFP-CB1R can then be synchronously released by addition of biotin and its trafficking through the secretory pathway and surface expression in both axons and dendrites can be monitored (Evans et al., 2017).

#### CB1R is directly trafficked to the axon through the secretory pathway

We first examined the synchronous trafficking of total SBP-EGFP-CB1R in the somatodendritic and axonal compartments of primary hippocampal neurons (**Fig. 1A-C**). Prior to biotin-mediated release, SBP-EGFP-CB1R was retained in the ER in the soma and dendrites but was absent from the axonal compartment and was not present at the cell surface (0 min; **Fig. 1A**). After addition of biotin, SBP-EGFP-CB1R moved through the secretory pathway and entered the axonal compartment at 25 min and continued to accumulate until 45 min when it reached its peak, which was comparable to an unretained control (O/N) (**Fig. 1B-C**). These data suggest that once released from the ER, CB1R is immediately trafficked towards the axonal compartment via the intracellular secretory pathway.

**Fig. 1.**
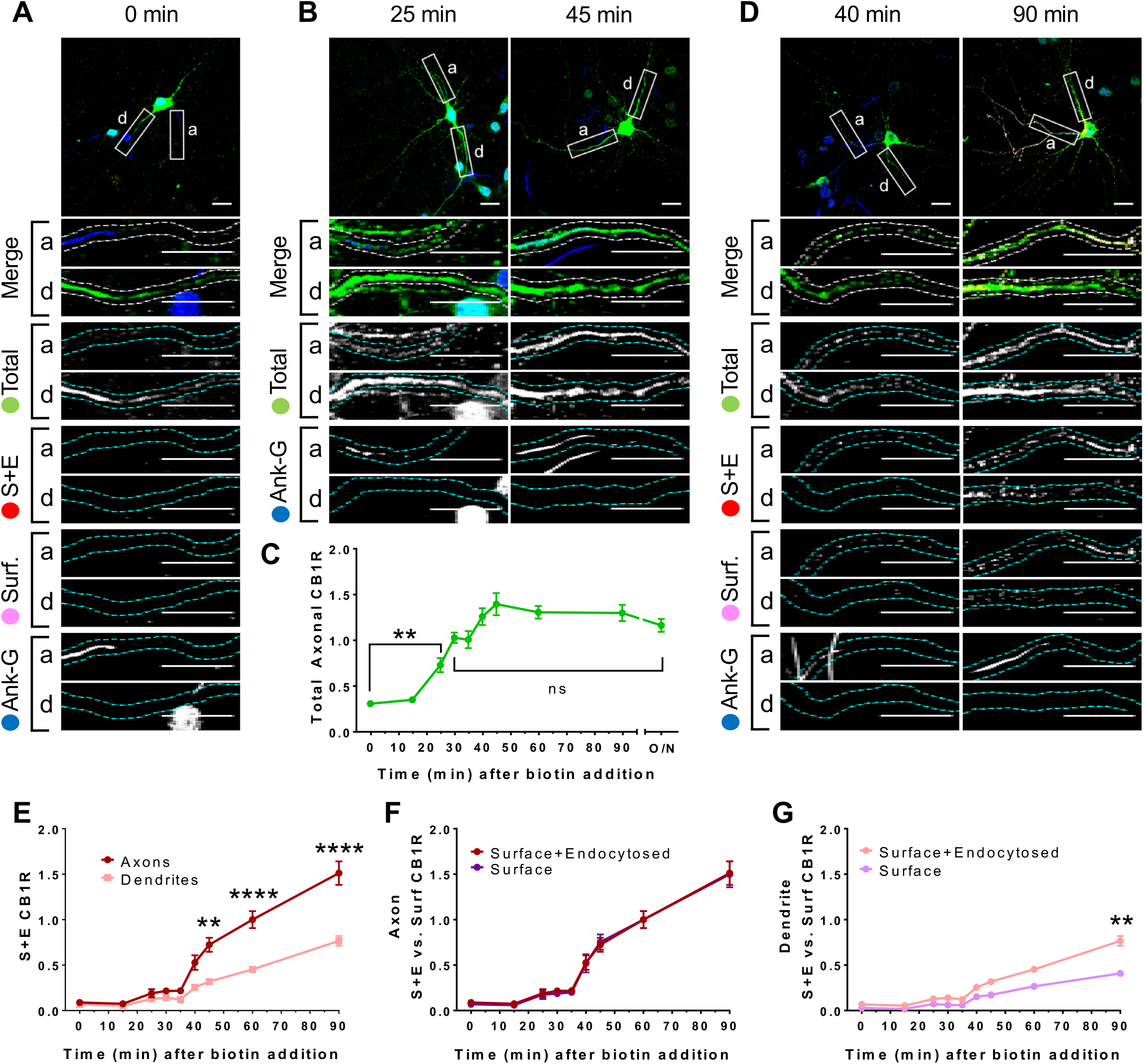
Newly synthesized CB1Rs are preferentially delivered to, and retained at, the axonal membrane to establish surface polarisation. The trafficking of SBP-EGFP-CB1R following release with biotin was monitored after 0 (no biotin), 15, 25, 30, 35, 40, 45, 60, 90 minutes and overnight (O/N; non-retained control) in DIV13 hippocampal neurons. Upper panels for each condition show whole cell field of view and lower panels are enlargements of axonal (a) and dendritic (d) ROIs. Green = total; red = surface+endocytosed; magenta = surface; blue = axon marker (Ankyrin-G). **A)** Representative image of a hippocampal neuron expressing RUSH SBP-EGFP-CB1R without biotin (0 min). SBP-EGFP-CB1R is anchored in the ER of the somatodendritic compartment and is not detected in the axonal compartment or on the surface of either compartment. Merge: green = total; blue = Ankyrin-G; red = surface+endocytosed; magenta = surface. **B)** Representative confocal images of hippocampal neurons expressing RUSH SBP-EGFP-CB1R 25 and 45 min after release showing that SBP-EGFP-CB1R has entered the axon. Merge: green = total; blue = Ankyrin-G. **C)** Quantification of data represented in (***A-B***). SBP-EGFP-CB1R was initially absent from the axon but entered after 25 minutes and continued to accumulate until it plateaued after 45 minutes to a level comparable to a non-retained control (O/N). One-way ANOVA with Tukey’s *post hoc* test. N = three to six independent experiments, n = 19-45 neurons per condition. (0 min vs. 25 min: 0.307±0.0173 vs. 0.729±0.0772; N = 6, n = 45 vs. N = 3, n = 19; **p = 0.0018. 30 min vs. ON: 1.03±0.0597 vs. 1.2±0.0632; N = 4, n = 32 vs. N = 4, n = 24, ^ns^p = 0.8186). **D**) Representative confocal images of DIV 13 hippocampal neurons expressing RUSH SBP-EGFP-CB1R 40 and 90 min after release showing that SBP-EGFP-CB1R is preferentially delivered to, and retained at, the axonal surface. Merge: surface to total seen as white; endocytosed to total seen as yellow. **E-G**) Two-way ANOVA with Tukey’s *post hoc* test (all analysed and corrected for multiple comparisons together). N = three to six independent experiments, n = 19-45 neurons per condition. **E)** Quantification of data represented in (***D***). SBP-EGFP-CB1R reached the surface of the axon 40 minutes after release and the surface of dendrites 60 minutes after release. Furthermore, significantly more SBP-EGFP-CB1R reached the axonal versus dendritic surface at 45, 60, and 90 minutes. (45 min, Axons vs. Dendrites: 0.723 ± 0.077 vs. 0.319 ± 0.035; N = 3, n = 20 vs. N = 3, n = 20; **p = 0.0054. 60 min, Axons vs. Dendrites: 1 ± 0.093 vs. 0.452 ± 0.023; N = 6, n = 46 vs. N = 6, n = 46; ****p < 0.0001. 90 min, Axons vs. Dendrites: 1.511 ± 0.129 vs. 0.566 ± 0.054; N = 4, n = 26 vs. N = 4, n = 26; ****p < 0.0001). **F)** Quantification of data represented in (***D***). Comparison between surface+endocytosed (red; see ***E***) and surface (magenta) curves show that SBP-EGFP-CB1R was retained on the surface of axons. (For all p > 0.9999). **G**) Quantification of data represented in (***D***). Comparison between surface+endocytosed (pale red; see ***E***) and surface (pale magenta) curves show that SBP-EGFP-CB1R was internalised from the surface of dendrites. (90 min, SE vs. S: 0.766 ± 0.054 vs. 0.408 ± 0.038; N = 4, n = 26 vs. N = 4, n = 26; **p = 0.0046).

#### *De novo* CB1R is more rapidly surface expressed in axons than in dendrites

Having established that SBP-EGFP-CB1R released from the ER traffics directly to axons, we next investigated where and when the newly synthesised SBP-EGFP-CB1R first reaches the plasma membrane. We determined how much SBP-EGFP-CB1R was surface expressed during a given time period using an antibody feeding assay (Evans et al., 2017). Antibody feeding was performed concurrent with the addition of biotin to release ER-retained SBP-EGFP-CB1R. This protocol labels both surface expressed CB1Rs and those that have been surface expressed and subsequently endocytosed (**Fig. 1D-G;** surface+endocytosed), giving a measure of total amount of surface expression irrespective of internalisation. SBP-EGFP-CB1R was surface expressed in axons 40 min after release from the ER, whereas in dendrites, CB1R was not surface expressed until 60 min after release (**Fig. 1E**). Moreover, significantly more SBP-EGFP-CB1R reached the surface of axons than the surface of dendrites 45, 60 and 90 min after release from the ER (**Fig. 1E**). These data demonstrate that the secretory pathway delivers a greater amount of CB1R more rapidly to the axonal membrane than to the dendritic membrane.

#### *De novo* CB1R is retained longer at the surface of axons than of dendrites

It has been suggested CB1R polarity is maintained by differential rates of endocytosis in the somatodendritic and axonal compartments (Leterrier et al., 2006; McDonald et al., 2007a). To test this, we also stained for surface SBP-EGFP-CB1R and compared the amount of surface expressed SBP-EGFP-CB1R to the amount of surface+endocytosed SBP-EGFP-CB1R in axons (**Fig. 1D,F**) and dendrites (**Fig. 1D,G**). In axons the normalised surface and surface+endocytosed curves were identical, suggesting that most surface expressed SBP-EGFP-CB1R is stable and retained at the membrane (**Fig. 1 D,F**). This may be due either to minimal endocytosis or to the efficient recycling of endocytosed receptors. In stark contrast, however, in dendrites there is significantly less surface than surface+endocytosed SBP-EGFP-CB1R 90 min after addition of biotin, indicating that surface expressed CB1R is more rapidly endocytosed from and/or not recycled back to the dendritic membrane (**Fig. 1G**).

Our results using RUSH time-resolved analysis show that CB1R surface polarity is established and maintained by two distinct but complementary mechanisms. Firstly, we show the novel finding that the secretory pathway preferentially delivers CB1R to the axonal surface, with significantly less going to the dendritic surface. Secondly, by distinguishing between surface and surface+endocytosed receptors, our antibody feeding experiments show that newly delivered CB1R is preferentially retained/stabilised at the axonal membrane and internalised from the dendritic membrane. Previous literature proposes that this differential internalisation is due to the presence of agonist in the dendritic membrane and absence of agonist on axonal membrane (Ladarre et al., 2014; Leterrier et al., 2006), although a potential role for constitutive internalisation distinct to agonist-induced internalisation has also been proposed (McDonald et al., 2007a). Taken together, we propose that preferential delivery and differential internalisation underpin the axonal surface polarisation of CB1R in hippocampal neurons.

### ctCB1R and *H9* can mediate axonal surface polarisation

While ctCB1R is implicated in desensitization and internalization (reviewed by (Mackie, 2008; Stadel et al., 2011)), the role of this region in determining axonal polarity has not been investigated. Furthermore, the function of the Helix 9 (*H9*) structural motif is unknown. We therefore wondered whether ctCB1R, or *H9* in particular, may contribute to CB1R surface polarisation.

To test this, we used CD4, a single-pass membrane protein that has no intrinsic localisation signals and is normally surface expressed in a non-polarised manner (Fache et al., 2004; Garrido et al., 2001). We expressed chimeras of CD4 alone, or CD4 fused to either ctCB1R^WT^ or a ctCB1R lacking the *H9* domain (ctCB1R^ΔH9^; **Fig. 2A**). In hippocampal neurons we examined each of the CD4 chimeras’ surface expression by immunostaining.

**Fig. 2.**
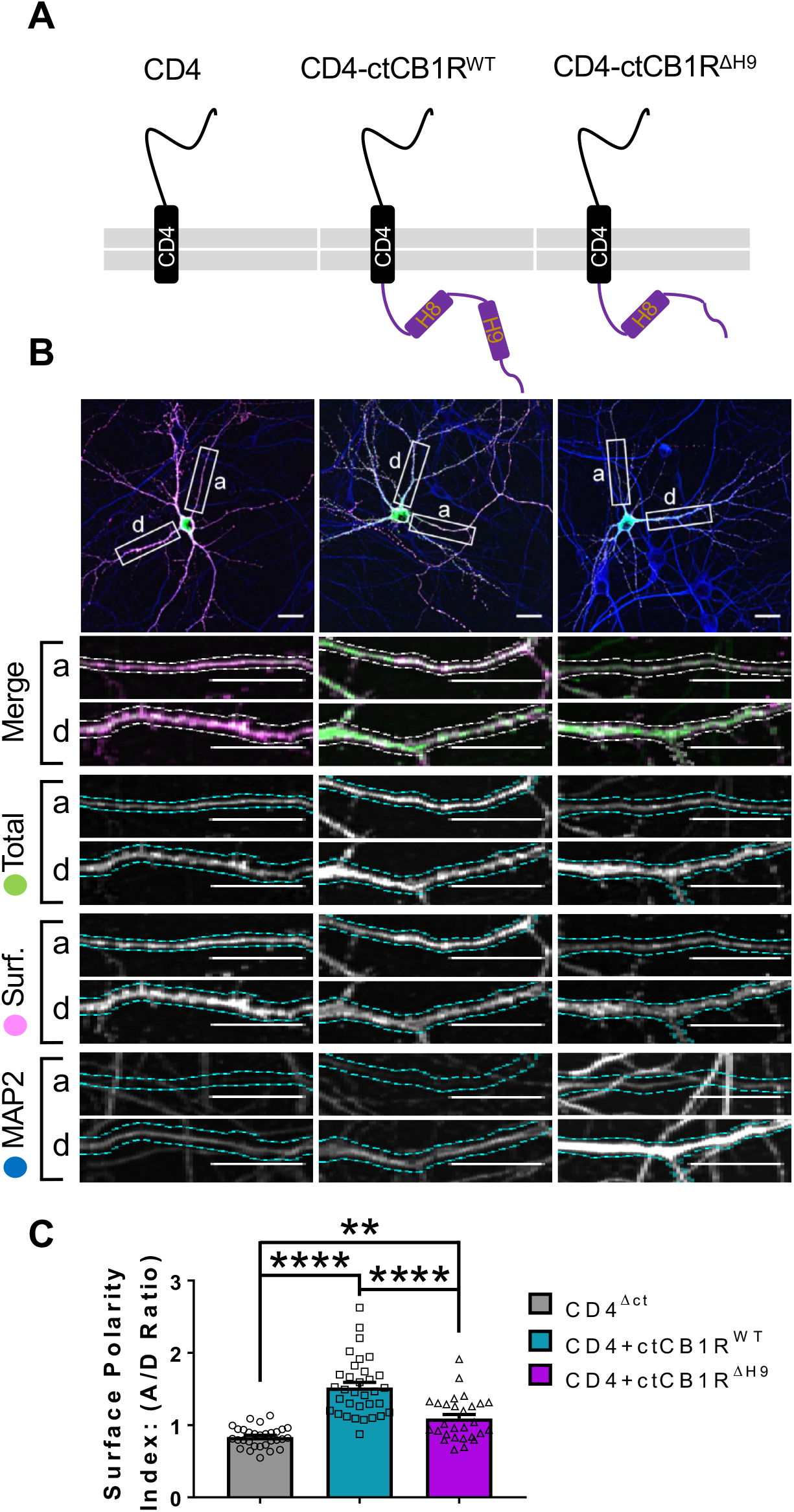
The C-terminal domain of CB1R, especially the Helix 9 motif, plays a role in axonal surface polarisation. **A**) Schematic of the CD4-ctCB1R chimeric proteins used in this Figure. **B**) Representative confocal images of hippocampal neurons showing the distribution of expressed CD4 (left), CD4-ctCB1R^WT^ (middle) or CD4-ctCB1R^ΔH9^ (right). Upper panels for each condition show a whole cell field of view and lower panels are enlargements of axonal (a) and dendritic (d) ROIs. Green = total; magenta = surface; blue = dendrite marker (MAP2). Merge: surface to total seen as white. **C**) Quantification of data represented in (***B***) presented as the surface polarity index (A/D ratio). CD4-ctCB1R^WT^ strongly favoured the axonal compartment compared to CD4 alone. CD4-ctCB1R^ΔH9^ favoured the axonal compartment significantly less than CD4-ctCB1R^WT^. One-way ANOVA with Tukey’s *post hoc* test. N = three independent experiments; n = 28-33 neurons per condition. (CD4 vs. WT: 0.834 ± 0.0255 vs. 1.52 ± 0.0696; N = 3, n = 30 vs. N = 3, n = 33; ****p <0.0001. CD4 vs. ΔH9: 0.834 ± 0.0255 vs. 1.09 ± 0.0562; N = 3, n = 30 vs. N = 3, n = 28; **p <0.0050. WT vs. ΔH9: 1.52 ± 0.0696 vs. 1.09 ± 0.0562; N = 3, n = 33 vs. N = 3, n = 28; ****p <0.0001).

Analysis of the axon to dendrite ratio of surface expression (the surface polarity index) revealed that CD4-ctCB1R^WT^ was markedly more axonally polarised than CD4 alone, indicating that ctCB1R may play a role in polarisation despite its lack of defined canonical localisation signals (**Fig. 2B, 2C**). Moreover, although still significantly axonally polarised, the degree of polarisation was significantly lower for CD4-ctCB1R^ΔH9^, suggesting that *H9* may also contribute to this process.

#### *H9* restricts delivery of CB1 R to the dendritic membrane

To further explore the possibility that *H9* is involved in the axonal surface polarity of CB1R, we used RUSH to compare the forward trafficking of SBP-EGFP-CB1R^WT^ and SBP-EGFP-CB1R^ΔH9^. As in **Fig. 1**, we labelled all the CB1R that had been surface expressed (surface+endocytosed) 0, 30, 60 and 90 min after biotin release from the ER (**Fig. 3A-G**).

**Fig. 3.**
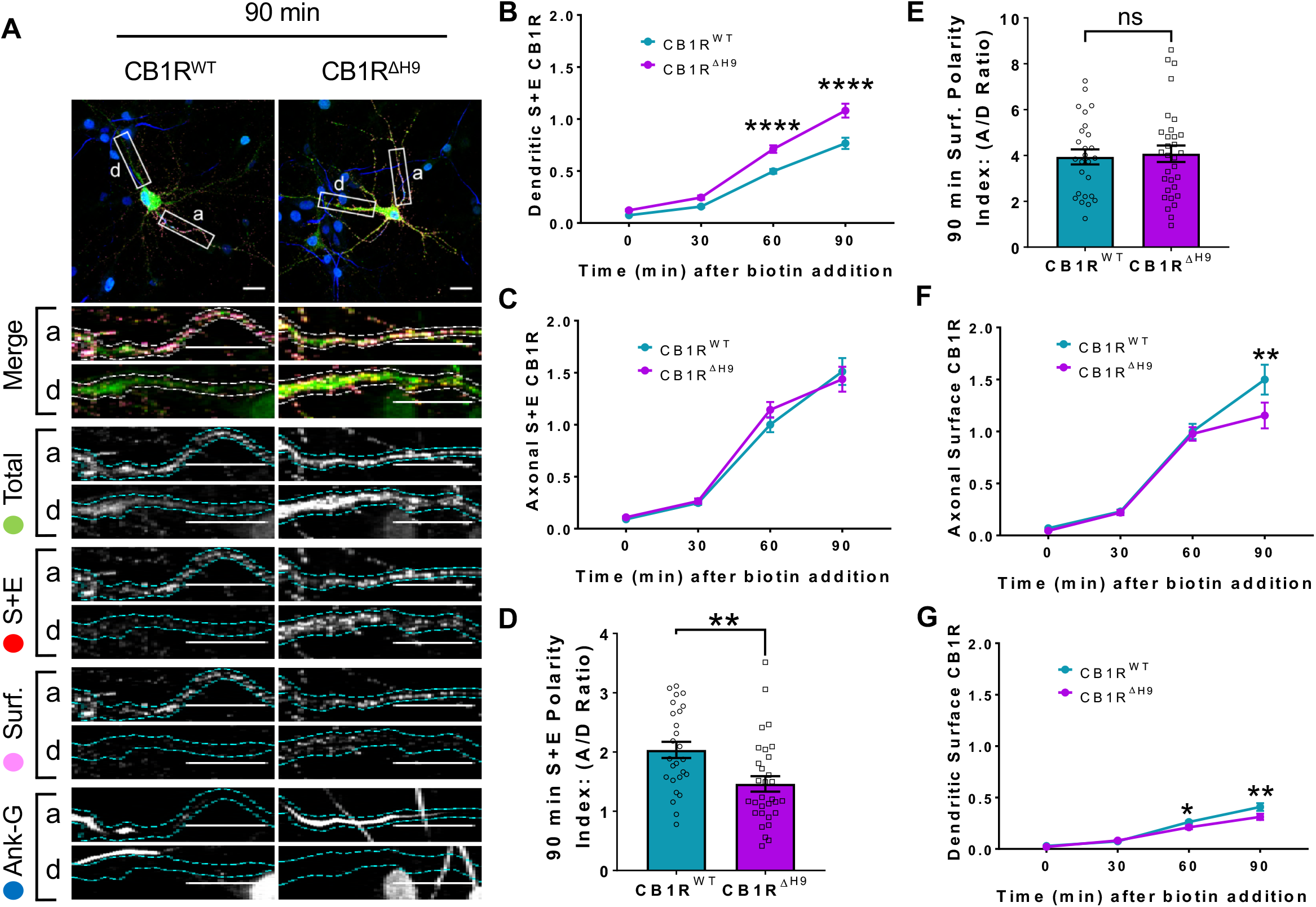
*H9* both restricts delivery of CB1R to the dendritic membrane and plays a role in surface retention of CB1R. The trafficking of RUSH SBP-EGFP-CB1R following release with biotin was monitored after 0 (no biotin), 30, 60, and 90 minutes in DIV 13 hippocampal neurons. **A**) Representative confocal images of hippocampal neurons expressing SBP-EGFP-CB1R^WT^ or SBP-EGFP-CB1R^ΔH9^ 90 minutes after release with biotin. Upper panels for each condition show whole cell field of view and lower panels are enlargements of axonal (a) and dendritic (d) ROIs. Green = total; red = surface+endocytosed; magenta = surface; blue = axon marker (Ankyrin-G). Merge: surface to total seen as white; endocytosed to total seen as yellow. **B**) Quantification of data represented in (***A***). Time-resolved analysis of surface+endocytosed receptors shows significantly more SBP-EGFP-CB1R^ΔH9^ reaches the surface of dendrites than SBP-EGFP-CB1R^WT^, indicating that *H9* may play a role in restricting delivery to the dendritic surface. Two-way ANOVA with Sidak’s *post hoc* test. Three to seven independent experiments, n = 26-63 neurons per condition. (60 min, WT vs. ΔH9: 0.497 ± 0.022 vs. 0.711 ± 0.036; N = 8, n = 63 vs. N = 8, n = 48; ****p <0.0001. 90 min, WT vs. ΔH9: 0.766 ± 0.054 vs. 1.08 ± 0.066; N = 4, n = 26 vs. N = 4, n = 31; ****p <0.0001). **C**) Quantification of data represented in (***A***). Time-resolved analysis of surface+endocytosed receptors shows no difference between SBP-EGFP-CB1R^WT^ and SBP-EGFP-CB1^ΔH9^ in reaching the surface of the axon. Two-way ANOVA with Sidak’s *post hoc* test. N = three to seven independent experiments, n = 26-63 neurons per condition. (0, 30, 60, 90 min; WT vs. ΔH9: p > 0.2459). **D**) Quantification of data represented in (***A***). Analysis of surface+endocytosed polarity demonstrates a defect in polarised delivery of SBP-EGFP-CB1R^ΔH9^ compared to SBP-EGFP-CB1R^WT^. Unpaired t-test. N = four independent experiments, n = 26-31 neurons per condition. (WT vs. ΔH9: 2.03 ± 0.136 vs. 1.46 ± 0.13; N = 4, n = 26 vs. N = 4, n = 31; **p = 0.0038). **E**) Quantification of data represented in (***A***). Analysis of surface polarity revealed no difference between SBP-EGFP-CB1R^WT^ and SBP-EGFP-CB1^ΔH9^. Unpaired t-test. N = four independent experiments, n = 26-31 neurons per condition. (3.935 ± 0.329 vs. 4.075 ± 0.361; N = 4, n = 26 vs. N = 4, n = 31; ^ns^p = 0.7797). **F**) Quantification of data represented in (***A***). Time-resolved analysis of surface receptors shows significantly less SBP-EGFP-CB1^ΔH9^ than SBP-EGFP-CB1R^WT^ on the surface of axons 90 minutes after release, most likely due to increased endocytosis of the Δ*H9* mutant. Two-way ANOVA with Sidak’s *post hoc* test. N = three to eight independent experiments, n = 26-63 neurons per condition. (90, WT vs. ΔH9: 1.498 vs. 1.154; N = 4, n = 26 vs. N = 4, n = 31; **p = 0.0066). **G**) Quantification of data represented in (***A***). Time-resolved analysis of surface receptors shows significantly less SBP-EGFP-CB1^ΔH9^ than SBP-EGFP-CB1R^WT^ on the surface of dendrites 60 and 90 minutes after release, most likely due to increased endocytosis. Two-way ANOVA with Sidak’s *post hoc* test. N = three to eight independent experiments, n = 26-63 neurons per condition. (60, WT vs. ΔH9: 0.262 ± 0.013 vs. 0.21 ± 0.018; N = 8, n = 63 vs. N = 8, n = 48; *p = 0.0232. 90, WT vs. ΔH9: 0.408 ± 0.038 vs. 0.312 ± 0.030; N = 4, n = 26 vs. N = 4, n = 31; **p = 0.0011).

Interestingly, significantly more SBP-EGFP-CB1R^ΔH9^ than SBP-EGFP-CB1R^WT^ reached the surface of dendrites during time course of our experiments (**Fig. 3B**), whereas trafficking to axons was similar for both SBP-EGFP-CB1R^WT^ and SBP-EGFP-CB1R^ΔH9^ (**Fig. 3C**). These altered properties resulted in a significant difference in the surface+endocytosed polarity index after 90 min (**Fig. 3D**) and are consistent with a role for *H9* in restricting delivery of CB1R to the dendritic membrane.

#### *H9* plays a role in the surface retention of CB1R

Surprisingly, however, in contrast to the total amount of CB1R that had been surface expressed during the time course (surface+endocytosed) (**Fig. 3D**) the polarity of the amount of CB1R on the cell surface 90 min after biotin-mediated release was identical for SBP-EGFP-CB1 R^WT^ and SBP-EGFP-CB1R^ΔH9^ (surface; **Fig. 3E**).

Closer analysis revealed identical levels of axonal surface expression of both SBP-EGFP-CB1R^WT^ and SBP-EGFP-CB1R^ΔH9^ 60 min after release from the ER. However, at 90 min there is significantly less surface expression of *ΔH9* mutant (**Fig. 3F**) suggesting that, although similar amounts of SBP-EGFP-CB1R^WT^ and SBP-EGFP-CB1R^ΔH9^ reach the surface, surface expression of SBP-EGFP-CB1R^ΔH9^ is less stable than that of the wild-type.

Furthermore, in dendrites, the increased delivery and surface trafficking of the *ΔH9* mutant is counteracted by the fact that less is retained at the surface 60 min after ER release (**Fig. 3G**).

Taken together these results suggest that, separate from its role in restricting delivery to the dendritic membrane, *H9* also plays a role in membrane stability and retention at both axons and dendrites.

### *H9* stabilises CB1R at the surface

To investigate the role of *H9* in membrane stability, we next compared surface expression (**Fig. 4A**) and endocytosis (**Fig. 4B**) of EGFP-CB1R^WT^ and EGFP-CB1R^ΔH9^ in axons and dendrites at steady-state. EGFP-CB1R^ΔH9^ displayed lower levels of surface expression (**Fig. 4C**), as well as increased endocytosis (**Fig. 4D**) in both axons and dendrites compared to EGFP-CB1R^WT^, suggesting *H9* plays a role in stabilising CB1R at the surface of both axons and dendrites. Moreover, similar to our findings using RUSH, there was there was no difference in surface polarity between wild-type and EGFP-CB1R^ΔH9^ (**Fig. 4E**). These findings suggest that, while *H9* plays a role in CB1R surface expression and endocytosis, its potential to mediate surface polarity is masked in the context of the full-length receptor.

**Fig. 4.**
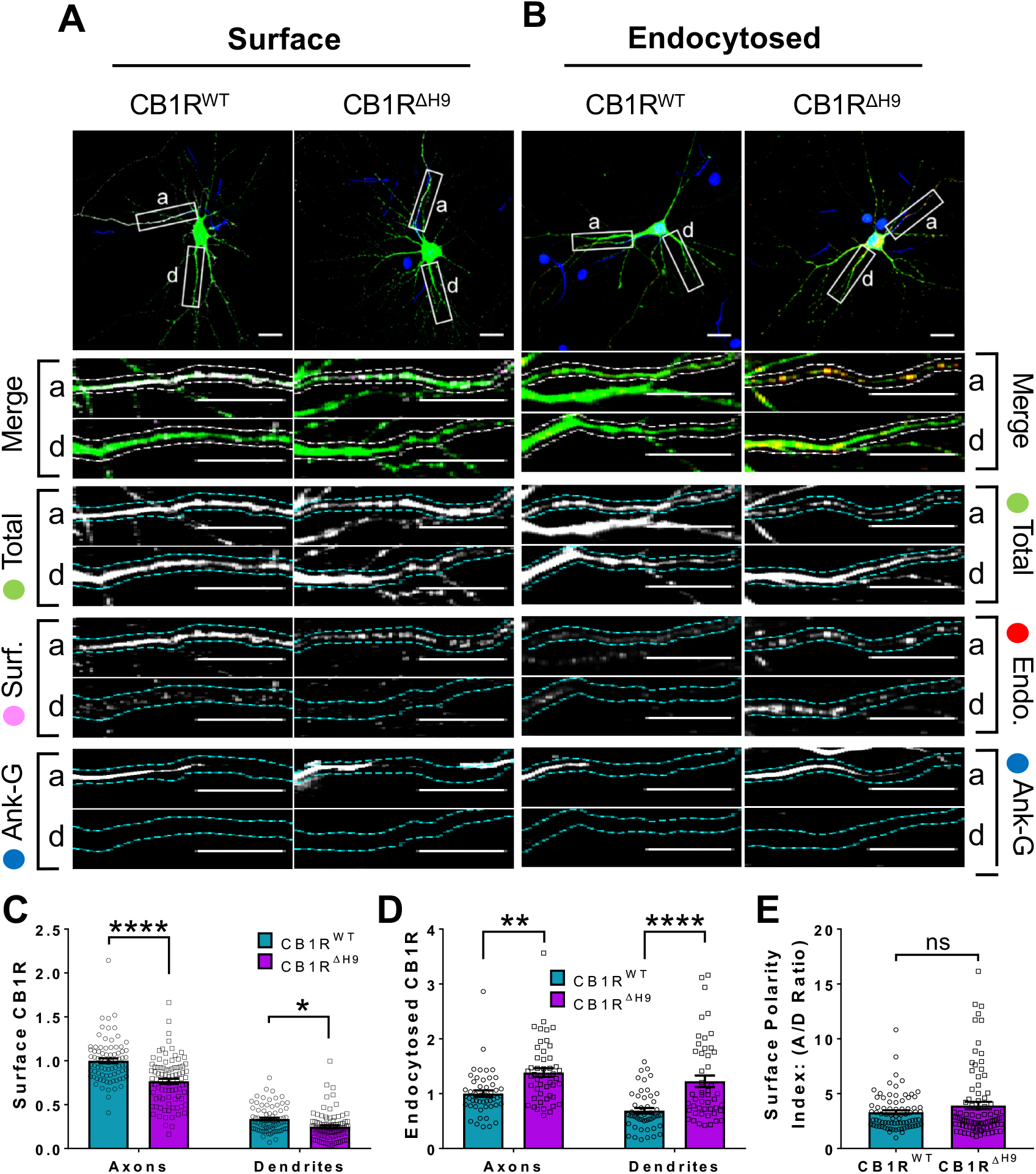
*H9* stabilises CB1R at the axonal surface. **A**) Representative confocal images of surface stained DIV 14 hippocampal neurons expressing EGFP-CB1R^WT^ or EGFP-CB1R^ΔH9^. Green = total; magenta = surface; blue = axon marker (Ankyrin-G). Merge: surface to total seen as white. **B**) Representative confocal images of DIV 14 primary hippocampal neurons expressing EGFP-CB1R^WT^ or EGFP-CB1R^ΔH9^. Neurons were subjected to 2 hours of antibody feeding followed by stripping off of surface antibody to reveal the endocytosed pool of receptors. Green = total; red = endocytosed; blue = axon marker (Ankyrin-G). Merge: endocytosed to total seen as yellow. **C**) Quantification of data shown in (A). Surface expression of EGFP-CB1R^ΔH9^ in both axons and dendrites was significantly reduced compared to EGFP-CB1R^WT^. Two-way ANOVA with Tukey’s *post hoc* test. N = ten independent experiments; n = 80-88 neurons per condition. (Axons, WT vs. ΔH9: 1 ± 0.028 vs. 0.765 ± 0.029; N = 10, n=80 vs. N = 10, n=88; ****p <0.0001. Dendrites, WT vs. ΔH9: 0.335 ± 0.016 vs. 0.247 ± 0.017; N = 10, n=80 vs. N = 10, n=88; *p = 0.0392). **D**) Quantification of data shown in (***B***). Endocytosis of EGFP-CB1R^ΔH9^ is significantly increased compared to EGFP-CB1R^WT^ in both axons and dendrites. One-way ANOVA with Tukey’s *post hoc* test. N = seven independent experiments; n = 49 neurons per condition. (Axons, WT vs. ΔH9: 1 ± 0.058 vs. 1.38 ± 0.08; **p = 0.0026. Dendrites, WT vs. ΔH9: 0.689 ± 0.05 vs. 1.225 ± 0.105; ****p <0.0001.) **E**) Quantification of data shown in (***A***) presented as the surface polarity index. There was no difference in surface polarity between EGFP-CB1R^WT^ or EGFP-CB1R^ΔH9^. Mann-Whitney test. N = ten independent experiments; n = 80-88 neurons per condition. (WT vs. ΔH9: 3.298 ± 0.1812 vs. 3.915 ± 0.3367; N = 10, n=80 vs. N = 10, n=88; p = 0.6886).

### CB1R^ΔH9^ is less efficient at activating downstream signalling pathways and more susceptible to agonist-induced internalisation

Because CB1R surface expression and polarisation has been linked to its activity (Ladarre et al., 2014; Leterrier et al., 2006), we next investigated if deleting *H9* affects CB1R downstream signalling pathways. We expressed EGFP-CB1R^WT^ or EGFP-CB1R^ΔH9^ in HEK293T cells, which contain no endogenous CB1R (Atwood et al., 2011), stimulated with the selective CB1R agonist ACEA (arachidonyl-2’-chloroethylamide) (Hillard et al., 1999) and monitored ERK1/2 phosphorylation as a measure of signalling downstream of CB1R (Daigle et al., 2008). There was no significant difference in ERK1/2 phosphorylation in cells expressing EGFP-CB1R^WT^ or EGFP-CB1R^ΔH9^ under basal conditions in the absence of ACEA. However, upon ACEA stimulation, the level of ERK1/2 activation was significantly reduced in EGFP-CB1R^ΔH9^-transfected cells compared to EGFP-CB1R^WT^-transfected cells expressing equivalent amounts of receptor (**Fig. 5A-C**), suggesting the *ΔH9* mutant is deficient in its ability to activate downstream signalling pathways.

**Fig. 5.**
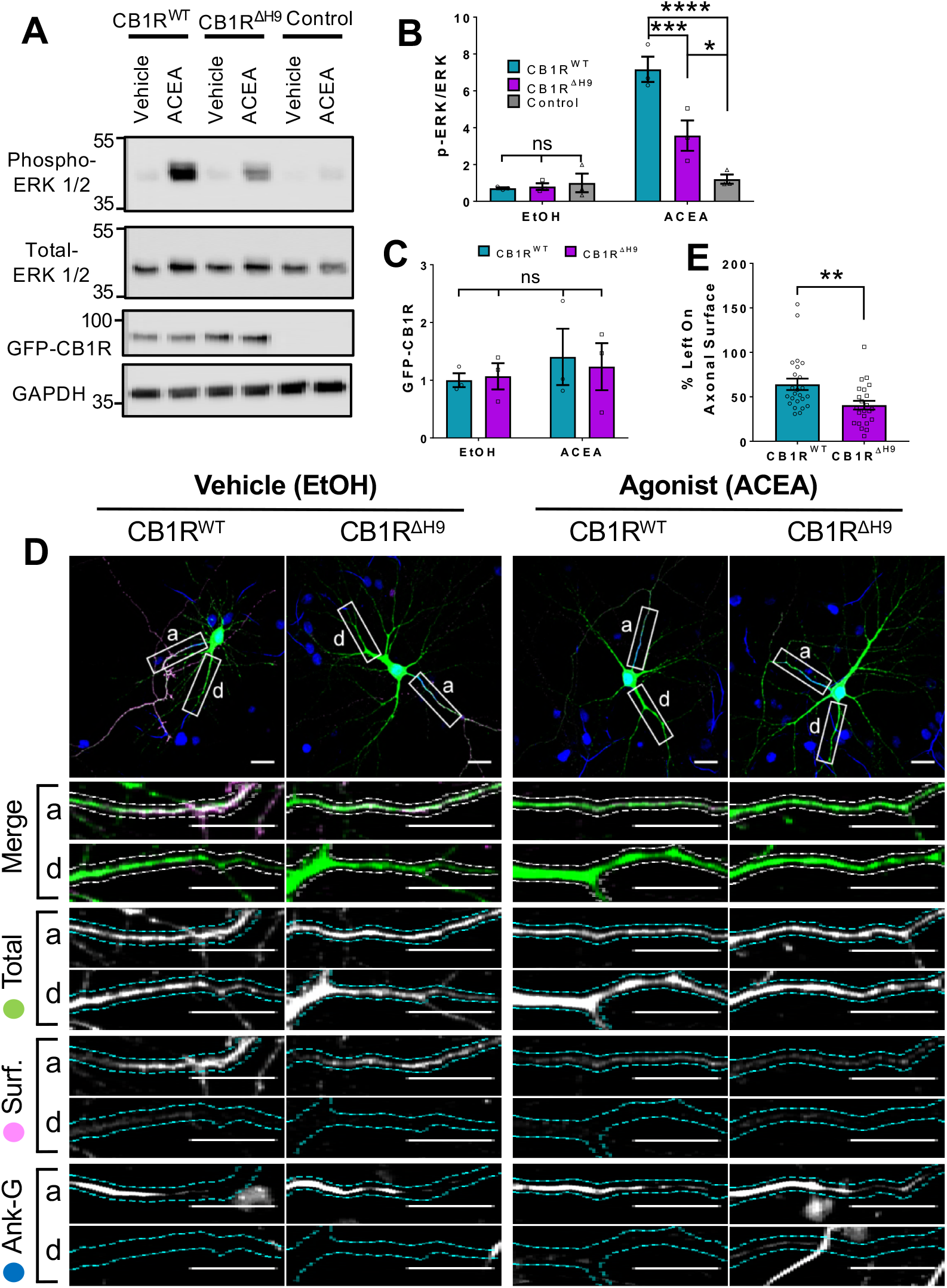
Role of *H9* in CB1R signalling and in resisting agonist-induced endocytosis. **A**) Representative blots showing ERK1/2 phosphorylation in HEK293T cells expressing EGFP-CB1R^WT^ or EGFP-CB1R^ΔH9^ following vehicle (0.1% EtOH) or ACEA (1μM) treatment for 5 minutes. **B**) Quantification of data shown in (***A***). Following treatment with ACEA, ERK1/2 was significantly more phosphorylated in EGFP-CB1R^WT^ - and EGFP-CB1R^ΔH9^-transfected cells compared to untransfected cells. However, ERK1/2 activation was significantly reduced in EGFP-CB1R^ΔH9^-expressing cells compared to EGFP-CB1R^WT^-expressing cells. There was no significant difference in ERK1/2 phosphorylation in vehicle-treated cells. Two-way ANOVA with Tukey’s *post hoc* test. N = three independent experiments. (ACEA, WT vs. Control: 7.17 ± 0.684 vs. 1.21 ± 0.252; ****p < 0.0001. ΔH9 vs. Control: 3.57 ± 0.825 vs. 1.21 ± 0.252; *p = 0.0150. WT vs. ΔH9: 7.17 ± 0.684 vs. 3.57 ± 0.825; ***p = 0.0007. EtOH, WT vs. ΔH9 vs. Control: ^ns^p ≥ 0.9125). **C**) Quantification of data shown in (***A***). EGFP-CB1R^WT^ and EGFP-CB1R^ΔH9^ expressed equally in HEK293T cells. Two-way ANOVA with Sidak’s *post hoc* test. Three independent experiments. (^ns^p ≥ 0.9654). **D**) Representative confocal images of DIV 12 hippocampal neurons expressing EGFP-CB1R^WT^ or EGFP-CB1R^ΔH9^ and treated with vehicle (0.1% EtOH) or CB1R agonist (5μM ACEA) for 3 hours. Upper panels for each condition show whole cell field of view and lower panels are enlargements of axonal (a) and dendritic (d) ROIs. Green = total; magenta = surface; blue = axon marker (Ankyrin-G). Merge: surface to total seen as white. **E**) Quantification of data represented in (***D***). Significantly less EGFP-CB1R^ΔH9^ than EGFP-CB1R^WT^ remained on the surface of axons after agonist application, indicating greater sensitivity to agonist-induced internalisation. The surface mean fluorescence was first normalised to the total mean fluorescence for each ROI, then to the average axonal EtOH value within a condition (set to 100%). Unpaired t-test. N = three independent experiments; n = 23-24 neurons per condition. (WT vs. ΔH9: 64 ± 6.42 vs. 40.6 ± 4.87; N = 3, n=24 vs. N = 3, n=23; **p = 0.0059).

We next monitored ACEA-induced internalisation of EGFP-CB1R^WT^ and EGFP-CB1R^ΔH9^ in axons of hippocampal neurons (**Fig. 5D**). ACEA-induced internalisation of EGFP-CB1R^ΔH9^ was significantly greater than that observed for EGFP-CB1R^WT^ (**Fig. 5E**). Taken together, these data indicate that CB1R^ΔH9^ is less stable at the axonal surface under basal conditions and that it is more susceptible to agonist-induced internalisation.

### The role of *H9* in polarity is revealed in the presence of inverse agonist

Our data thus far have indicated that ctCB1R, and the *H9* domain in particular, can mediate surface polarity of a CD4 chimera (**Fig. 2**), and promote polarised surface delivery of CB1R (**Fig. 3**). In contrast, deletion of *H9* has no effect on CB1R surface polarity at steady-state (**Fig. 4**). However, deletion of *H9* does have a striking effect on the surface stability of CB1R – CB1R^ΔH9^ is less surface expressed in both axons and dendrites and shows increased endocytosis (**Figs. 3 and 4**). Furthermore, CB1R ^ΔH9^ is more responsive to agonist-induced internalisation (**Fig. 5**). We therefore wondered whether the difference between the CD4 chimeras and the full-length receptor, and the difference between surface+endocytosed and surface polarity, may be due to the agonist binding capability of the full-length receptor. Inverse agonist treatment, which prevents the receptor entering an active conformation, has previously been shown to increase somatodendritic surface expression similarly to treatment with an endocytosis inhibitor (Leterrier et al., 2006). We thus reasoned that in this case, inverse agonist treatment may reveal a difference in surface polarity between EGFP-CB1R^WT^ and EGFP-CB1R^ΔH9^, like that observed with the CD4 chimeras and in surface+endocytosed polarity.

We treated hippocampal neurons expressing either EGFP-CB1R^WT^ or EGFP-CB1R^ΔH9^ with the CB1R-specific inverse agonist AM281 (Leterrier et al., 2004) (**Fig. 6A**). In the DMSO control both EGFP-CB1R^WT^ and EGFP-CB1R^ΔH9^ displayed similar levels of surface polarity. In the presence of AM281, however, EGFP-CB1R^ΔH9^ had significantly reduced surface polarity compared EGFP-CB1R^WT^ (**Fig. 6B**) due to a significantly increased amount of dendritic surface expression (**Fig. 6C**).

**Fig. 6.**
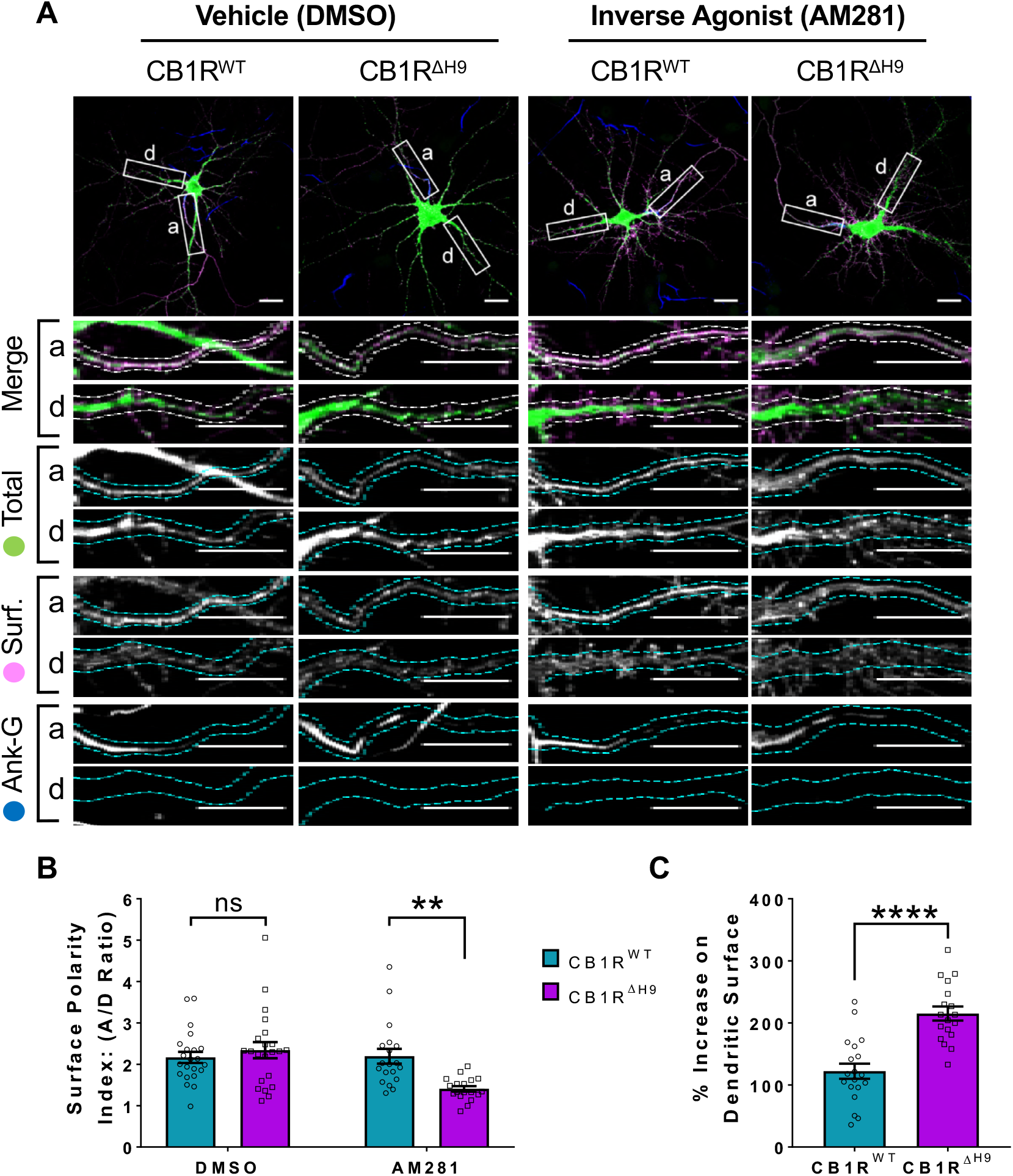
The role of *H9* in polarity is revealed in the presence of inverse agonist. **A**) Representative confocal images of DIV 14 hippocampal neurons expressing EGFP-CB1R^WT^ or EGFP-CB1R^ΔH9^ and treated with vehicle (0.2% DMSO) or CB1R inverse agonist (10μM AM281) for 3 hours. Upper panels for each condition show whole cell field of view and lower panels are enlargements of axonal (a) and dendritic (d) ROIs. Green = total; magenta = surface; blue = axon marker (Ankyrin-G). Merge: surface to total seen as white. **B**) Quantification of data shown in (***A***) presented as the surface polarity index (A/D ratio). In the presence of inverse agonist, but not vehicle, EGFP-CB1R^ΔH9^ was significantly less axonally polarised than EGFP-CB1R^WT^. Two-way ANOVA with Sidak’s *post hoc* test. N = three independent experiments; n = 18-22 neurons per condition. (DMSO, WT vs. ΔH9: 2.17 ± 0.135 vs. 2.34 ± 0.196; N = 3, n = 22 vs. N = 3, n = 22; ^ns^p = 0.9605. AM281, WT vs. ΔH9: 2.2 ± 0.18 vs. 1.41 ± 0.0649; N = 3, n = 19 vs. N = 3, n = 18; **p = 0.0067). **C**) Quantification of data represented in (***A***). Significantly more EGFP-CB1R^ΔH9^ than EGFP-CB1R^WT^ relocated to the surface of dendrites after inverse agonist application. The surface mean fluorescence was first normalised to the total mean fluorescence for each ROI, then to the average DMSO value within a condition (set to 100%). Unpaired t-test. N = three independent experiments; n = 18-19 neurons per condition. (WT vs. ΔH9: 122 ± 12.2 vs. 215 ± 11.3; N = 3, n = 19 vs. N = 3, n = 18; ****p < 0.0001).

These data suggest that in the absence of constitutive activity of the receptor, *H9* plays a role in mediating CB1R surface polarity. Furthermore, these data suggest that the increased internalisation observed in dendrites with *H9* deletion may be mediated by the presence of agonist. Finally, our findings reaffirm the importance the state-dependent effect on CB1R trafficking.

## Discussion

Our data indicate that axonal surface polarity of CB1R occurs as a result of two distinct, but complementary, mechanisms. *1)* Using time-resolved RUSH assays we demonstrate that more *de novo* CB1R is delivered to the axon and that it is more rapidly surface expressed than in dendrites. *2)* Once at the axonal membrane the newly delivered CB1R is more stably retained whereas in dendritic membrane CB1R surface expression is transient and it is rapidly internalised. However, we also note that our data do not specifically exclude the possibility that CB1R internalised into the somatodendritic endocytosed compartment can be rerouted to the axon via the transcytosis pathway, thus further facilitating axonal polarity (Simon et al., 2013).

Furthermore, since CD4-ctCB1R^WT^ and CD4-ctCB1R^ΔH9^chimeras cannot bind agonist, our results are consistent with ctCB1R contributing to constitutive polarisation via a mechanism distinct from the proposed continuous activation of CB1R by the presence of the endogenous agonist 2-Arachidonoylglycerol (2-AG) in the dendritic membrane (Ladarre et al., 2014). Our data suggest that ctCB1R, especially *H9*, plays a role in constitutive preferential delivery of CB1R to the axonal membrane.

Our results further demonstrate that ctCB1R is important for maintaining axonal surface polarity, in part mediated by the *H9* motif, which plays a role in both the preferential delivery and selective retention of CB1R at in axons. We show that deleting *H9* (CB1R^ΔH9^) has a range of effects on trafficking, surface expression and signalling in hippocampal neurons. More specifically, these include; *i)* CB1R^ΔH9^ lacks the preferential delivery to axons observed for CB1R^WT^, *ii*) CB1R^ΔH9^ is less efficiently surface expressed, *iii*) CB1R^ΔH9^ that does reach the surface it is more rapidly endocytosed in both axons and dendrites and *iv*) CB1R^ΔH9^ is more sensitive to agonist-induced internalisation and less efficient at downstream signalling, monitored by activation of ERK1/2 phosphorylation.

### Preferential axonal trafficking

The mechanism behind polarised membrane trafficking in neurons is a fundamental question and our data suggest a sorting mechanism at the level of the secretory pathway that preferentially targets CB1R to the axon. Since dendritic and axonal cargo are synthesized in the somatodendritic compartment, selective sorting to the correct domain is crucial. While several sorting signals and adaptors have been described for dendritic cargo, the mechanisms behind selective sorting to axons are less well known (Lasiecka and Winckler, 2011, Bentley, 2016 #43663). For example, a recent study in *C. elegans* has suggested that sorting of cargos to axons or dendrites depends on binding to different types of clathrin-associated adaptor proteins (AP); axonal cargo bind to AP-3 whereas dendritic cargo bind to AP-1 (Li et al., 2016). Interestingly, AP-3 binding has been associated with CB1R trafficking to the lysosome in the dendritic compartment (Rozenfeld and Devi, 2008). One possibility is that *H9* may modulate CB1R binding to AP-3, reducing both preferential delivery to axons and, perhaps, reducing sorting to lysosomes, causing an increase in dendritic membrane CB1R. More studies are needed to examine the possibility of *H9* influencing AP-3 and CB1R interaction.

### *H9* and membrane retention

Our data suggest that *H9* stabilises CB1R at the membrane, regardless of compartment. While the *H8* domain is highly conserved in GPCRs, structural domains analogous to *H9* have only been reported in squid rhodopsin (Murakami and Kouyama, 2008) and the bradykinin receptor (Piserchio et al., 2005). NMR and circular dichroism studies suggest that *H9*, like *H8*, is an amphipathic α-helix, associating with the lipid bilayer via a cluster of hydrophobic residues on the non-polar face of the helix (Ahn et al., 2009). Furthermore, *H9* contains a cysteine residue, raising the possibility that posttranslational modifications such as palmitoylation, prenylation or farnesylation could modulate membrane association (Tortosa and Hoogenraad, 2018).

Since our data suggest that *H9* stabilises CB1R at the membrane, it is possible that the membrane association of *H9* could mask internalisation signals or interacting motifs. Consistent with this possibility, ctCB1R interacting proteins regulate CB1R endocytosis. SGIP1, a protein linked to clathrin-mediated endocytosis, prevents internalisation of activated CB1R (Hajkova et al., 2016). Similarly, cannabinoid receptor interacting protein 1a (CRIP1a) reduces constitutive CB1R internalisation (Mascia et al., 2017) by competing with β-Arrestin binding (Blume et al., 2017).

Therefore, it is possible that *H9* mediates the interactions between CB1R and SGIP1 and/or selectively promotes β-Arrestin rather than CRIP1a binding. Further studies examining the interaction between CB1R^WT^, CB1R^ΔH9^, CRIP1a, β-Arrestin1/2, and SGIP1 are needed to examine the mechanism by which *H9* stabilises surface CB1R.

Given the increased interest in CB1R as a clinical target, understanding the fundamental cell biology and trafficking behaviour of CB1R is an increasingly active and important area of research. Taken together, our results reveal that the C-terminal domain, and *H9* in particular, play important roles in trafficking of CB1R. These findings provide important insight into the mechanisms of CB1R polarity and highlight *H9* as an important regulator of CB1R endocytosis and surface expression.

## Materials and Methods

### Constructs and reagents

A rat CB1R construct lacking residues 1-25, containing the putative mitochondrial targeting sequence (Hebert-Chatelain et al., 2016), was used as a template for sub-cloning into pcDNA3.1 (McDonald et al., 2007b). Helix 9 (residues 440-460) was removed by site-directed mutagenesis. These WT and ΔH9 constructs were subsequently used as a template to clone into the RUSH vector system (interleukin-2 signal peptide followed by SBP and EGFP N-terminal tags) as previously described (Boncompain and Perez, 2013; Evans et al., 2017). Non-ER-retained SBP-EGFP-tagged versions were obtained by re-cloning these inserts from the RUSH vector into pcDNA3.1 (SS_Ile2_-SBP-EGFP-CB1R). Chimeric CD4-ctCB1R WT and Δ*H9* were generated by overlap extension PCR followed by cloning into a plasmid expressing CD4 lacking its own C-terminus (Garrido et al., 2001).

Chicken anti-GFP was from Abcam (ab13970); mouse anti-Ankyrin-G was from NeuroMab (clone N106/36); rabbit anti-MAP2 was from Synaptic Systems (188 003); mouse anti-CD4 was from BioLegend (clone OKT4); rat anti-GFP was from ChromoTek (3H9); anti-phosphoERK (M7802), and anti-non-phosphoERK (M3807) were from Sigma; mouse anti-GAPDH (6C5 ab8245) was from Abcam. All fluorescent secondaries were from Jackson Immunoresearch Laboratories and HRP conjugated secondaries were from Sigma. ACEA and AM281 were from Tocris bio-techne.

### Cell culture and Transfection

Dissociated hippocampal cultures were prepared from E17-E18 Wistar rats as previously described (Martin and Henley, 2004). Glass coverslips were coated in poly-D-lysine or poly-L-lysine (1mg/mL, Sigma) in borate buffer (10mM borax, 50mM boric acid) overnight and washed in water. Dissociated hippocampal cells were plated at different densities in plating medium (Neurobasal, Gibco supplemented with 10% horse serum, Sigma; 2 mM GlutaMAX, Gibco; and either GS21, GlobalStem, or B27, Thermo Fisher) which was changed to feeding medium (Neurobasal supplemented with 1.2 mM GlutaMAX and GS21 or B27) after 24 hours. For RUSH experiments, cells were plated and fed in media containing GS21 instead of B27 because it does not contain biotin. Cells were incubated at 37°C and 5% CO2 for up to 2 weeks. Animal care and procedures were carried out in accordance with UK Home Office and University of Bristol guidelines.

Transfection of neuronal cultures was carried out at DIV 12 using Lipofectamine2000 (Invitrogen) according to the manufacturer’s instructions with minor modifications. Cells were left for 20-48 hours before fixation.

### Phospho-ERK assay

HEK293T cells were transfected with EGFP-CB1R^WT^, EGFP-CB1R^ΔH9^, or empty pcDNA3.1 and left for 24 hours. The cells were serum-starved overnight and then treated with 1μM ACEA or 0.01% EtOH for 5 min before being lysed in lysis buffer (50mM Tris-HCl; 150mM NaCl; 1% CHAPS, ThermoFisher Scientific; protease inhibitors, Roche) with phosphatase inhibitors (Pierce, ThermoFisher Scientific). SDS-PAGE and Western blotting procedures were carried out according to standard protocols.

### Live surface staining and antibody feeding

To measure surface staining, cultured neurons were cooled at room temperature for 5-10 min, then incubated with the appropriate antibody (chicken anti-GFP or mouse anti-CD4) in conditioned media for 10-20 min at RT. The neurons were washed multiple times in PBS before fixation.

For agonist and inverse agonist experiments, the neurons were treated with 5μM ACEA (in EtOH) or vehicle control (0.1% EtOH) for 3 hours or 10μM AM281 (in DMSO) or vehicle control (0.2% DMSO) for 3 hours in conditioned media at 37°C and 5% CO2, and then subsequently surface stained.

To measure endocytosed receptors, neurons were fed with chicken anti-GFP for 2h in conditioned media at 37°C and 5% CO2. Neurons were washed several times in PBS and then surface antibody was stripped by 2 quick washes with ice-cold pH 2.5 PBS followed by several washes in PBS before fixation.

### RUSH live labelling

Neurons were transfected with RUSH constructs at DIV 12 for no longer than 24 hours to prevent ER stress resulting from accumulation of unreleased receptors. Neurons were incubated in conditioned media containing D-biotin (40μM, Sigma) and chicken anti-GFP (1:1,000) for different lengths of time at 37°C and 5% CO2. The 0 min timepoint was only incubated with chicken anti-GFP without biotin for 60 min. For the O/N timepoint, neurons were incubated in 40μM D-biotin immediately following transfection and then left overnight at 37°C and 5% CO2 before being incubated with biotin and chicken anti-GFP for 60 min to label surface CB1R. Every independent experiment included a 60 min timepoint to which values were normalised and a 0 min control. Following biotin treatment, neurons were washed several times in PBS and cooled to 4°C to prevent further internalisation. They were then live labelled with 647-labelled anti-chicken in conditioned media for 15 min at 4°C before being fixed and permeabilised and stained with Cy3-labelled anti-chicken. In the text, “surface” thus refers to 647 fluorescence acquisition, whereas “surface+endocytosed” refers to Cy3 fluorescence acquisition.

### Fixation and fixed immunostaining

Cultured neurons were fixed in 4% formaldehyde in PBS for 12 min, then washed 3x in PBS, 1x in 100mM Glycine in PBS, and 3x in PBS. The neurons were then blocked and permeabilised in PBS + 3% BSA + 0.1% Triton X-100 before being incubated in fluorescent secondary (1:400) in PBS + 3% BSA. Subsequently, the neurons were re-incubated in primary antibody (anti-GFP or anti-CD4) to measure total levels of expression and stained with either anti-MAP2 (dendritic marker) or anti-Ankyrin-G (axonal initial segment marker) in PBS + 3% BSA. The neurons were then washed several times in PBS and mounted onto glass slides using Fluoromount-G (ThermoFisher Scientific).

### Image acquisition and analysis

Images were acquired using either a Leica SPE single channel confocal laser scanning microscope or a Leica SP8 AOBS confocal laser scanning microscope (Wolfson Bioimaging Facility, University of Bristol). All settings were kept the same within experiments. Neurons used for data acquisition were selected only on their total staining.

All quantification was performed using ImageJ software. Based on previous experiments, at least five cells were analysed per experiment, and at least three independent experiments (i.e. on different neuronal cultures on different days) were performed.

Images were max projected, and regions of interest (ROIs) of approximately similar lengths were drawn around axons and 3-4 proximal and secondary dendrites based on the total channel only. Axons were defined either as processes whose initial segment was positive for Ankyrin-G or as processes negative for MAP2. The mean fluorescence was measured for each channel and the dendritic values were averaged. “Surface” or “endocytosed” mean fluorescence values were normalised to the “total” mean fluorescence value for each ROI to account for varying levels of expression of transfected constructs. These values were then normalised to the axon value of the control (WT, WT + vehicle, or CD4).

Because of the change in total mean fluorescence in axons throughout the different conditions, the above image analysis was slightly modified for RUSH experiments. In these experiments, neurites were traced using NeuronJ so that only the mean fluorescence of exactly the first 50μm of the axons and 30-40μm of 2-4 primary dendrites for each channel was measured. All “surface” and “surface+endocytosed” values (of both axons and dendrites) were normalised to the average total dendritic value for each neuron. Axon total mean fluorescence was also normalised to the average total dendritic value within each cell. All values were then normalised to the WT 60 min axon value within each experiment.

“Polarity indices (A/D ratio)” were calculated by dividing the axonal mean fluorescence value by the average dendritic mean fluorescence value.

The scalebar for all images represents 20μm.

### Statistics

All statistics were performed using GraphPad Prism. The ROUT method was used to identify outliers for all parameters measured before normalising to control. Neurons were removed from analysis if any one parameter was found to be an outlier. To determine statistical significance between two groups, a D’Agostino & Pearson normality test was performed. Unpaired t-tests were performed on data that passed the normality test whereas the Mann-Whitney test was used if it did not. One- or Two-way ANOVAs with Tukey’s or Sidak’s *post hoc* test were used to determine statistical significance between more than two groups depending on the comparisons required. *p ≤ 0.05, **p ≤ 0.01, ***p ≤ 0.001, ****p ≤ 0.0001. All data are presented as mean ± SEM.

## Author Contributions

AFJ and KLH performed the experiments. AJE, YN and KAW provided constructs, technical assistance and advice. JMH managed the project. All authors contributed to writing and editing the manuscript.

## Acknowledgements

We are grateful to the MRC, BBSRC, Wellcome Trust and ERC for financial support. AFJ is funded by a University of Bristol PhD Scholarship. KLH was funded by an MRC PhD studentship. AJE was funded by a Wellcome Trust PhD studentship. We thank F. Perez and G. Boncompain (Institut Curie, Paris) for the RUSH constructs, A. Irvine (University of Dundee) for CB1R plasmids and B. Dargent (Universite de la Mediterranee, Marseille) for CD4 plasmids. We also thank the Wolfson Bioimaging Facility at the University of Bristol.

